# Potent and pan-neutralization of SARS-CoV-2 variants of concern by DARPins

**DOI:** 10.1101/2022.05.30.493765

**Authors:** Vikas Chonira, Young Do Kwon, Jason Gorman, James Brett Case, Zhiqiang Ku, Rudo Simeon, Ryan G. Casner, Darcy R. Harris, Adam S. Olia, Tyler Stephens, Lawrence Shapiro, Hannah Boyd, Yaroslav Tsybovsky, Florian Krammer, Michael S. Diamond, Peter D. Kwong, Zhiqiang An, Zhilei Chen

**Author notes:** These authors contributed equally (Young Do Kwon, Jason Gorman, James Brett Case, Zhiqiang Ku).

## Abstract

We report the engineering and selection of two synthetic proteins – FSR16m and FSR22 – for possible treatment of SARS-CoV-2 infection. FSR16m and FSR22 are trimeric proteins composed of DARPin SR16m or SR22 fused with a T4 foldon and exhibit broad spectrum neutralization of SARS-Cov-2 strains. The IC_50_ values of FSR16m against authentic B.1.351, B.1.617.2 and BA.1.1 variants are 3.4 ng/mL, 2.2 ng/mL and 7.4 ng/mL, respectively, comparable to currently used therapeutic antibodies. Despite the use of the spike protein from a now historical wild-type virus for design, FSR16m and FSR22 both exhibit increased neutralization against newly-emerged variants of concern (39- to 296-fold) in pseudovirus assays. Cryo-EM structures revealed that these DARPins recognize a region of the receptor binding domain (RBD, residues 455-456, 486-489) overlapping a critical portion of the ACE2-binding surface. K18-hACE2 transgenic mice inoculated with a B.1.617.2 variant and receiving intranasally-administered FSR16m were protected as judged by less weight loss and 10-100-fold reductions in viral burden in the upper and lower respiratory tracts. The strong and broad neutralization potency make FSR16m and FSR22 promising candidates for prevention and treatment of infection by current and potential future strains of SARS-CoV-2.

## INTRODUCTION

Severe acute respiratory syndrome coronavirus 2 (SARS-CoV-2) has infected over 518 million people worldwide resulting in over 6.2 million deaths as of May 2022^1^. Multiple SARS-CoV-2 variants with increased infectivity have emerged, which jeopardize the utility of current vaccines and therapeutic antibodies. Although several monoclonal antibody (mAb) therapeutics have received emergency use authorization and demonstrated efficacy in patients, they are limited by high production cost, global supply issues, and inconvenient routes of administration^2^. In addition, many human-derived mAbs show reduced efficacy against newly-evolved viral variants^3^.

Here, we report the engineering and characterization of two highly potent and broadly neutralizing synthetic proteins, FSR16m and FSR22, as candidates for preventing and treating coronavirus disease 2019 (COVID-19). The active domains of FSR16m and FSR22 are designed ankyrin repeat proteins (DARPins) engineered to mimic human angiotensin converting enzyme 2 (hACE2) binding to the receptor binding domain (RBD) of the spike protein. DARPin is a versatile synthetic binder scaffold that has high thermostability and has been engineered to bind an array of targets with pico- to nanomolar affinities^4,5^, including the SARS-CoV-2 spike protein^6,7^. Trimerization of RBD-binding DARPins SR16m and SR22 with a T4 foldon^8^ molecule increased their neutralization potency by 35,000- and 3,800-fold, respectively. Remarkably, and despite using the Wuhan-1 historical isolate spike protein as the target of engineering, both FSR16m and FSR22 exhibit 39- to 296-fold enhanced neutralization potency against a panel of SARS-CoV-2 variants of interest (VOI) and concern (VOC) relative to the ancestral virus in the pseudovirus assay. Cryo-electron microscopy (cryo-EM) studies confirmed that both SR16m and SR22 target an essential ACE2-binding epitope on the RBD, providing support for the idea that these DARPins can remain effective against future SARS-CoV-2 variants. Although the improvement in neutralizing potency against some newly-emerged viral variants was not built into our DARPins design, the broad-spectrum activity is a direct result of our engineering approach that combines protein trimerization to increase avidity with the selection of DARPins that can compete with the viral receptor (*i*.*e*., hACE2).

## RESULTS

### Engineering of FSR16m and FSR22

Employing an in-house DARPin library with >10^9^ distinct clones displayed on M13 phage particles, we sequentially enriched binders to biotinylated RBD and the full-length spike protein (Wuhan-1 strain) over four rounds of phage panning (**Figure S1**). The enriched DARPin library pool from the final round was cloned into an expression vector with 6xHis and Myc tags at the N-terminus and transformed into BL21(DE3) *E. coli* cells. Three hundred colonies were picked and grown in 96-well plates, and the cell lysate was subjected to enzyme-linked immunosorbent assay (ELISA)-based screens. Eleven unique clones bound strongly to both the RBD and the full-length (fl-) spike protein and were purified by nickel-affinity chromatography. A competition ELISA was used to identify DARPin molecules that could inhibit binding between hACE2 and spike proteins, which yielded two unique clones: SR16 and SR22. Sodium dodecyl sulphate-polyacrylamide gel electrophoresis (SDS-PAGE) analysis of these proteins revealed a homogenous band for SR22 but an additional band for SR16 (**Figure S2**). The N-terminal 6xHis tag in SR16 was moved to the C-terminus to give rise to SR16m.

Since the SARS-CoV-2 spike protein adopts a trimeric structure, we trimerized both SR16m and SR22 through fusion to a T4 foldon^8^, a small trimeric protein, with a flexible (GGGGSLQ)x2 linker to form FSR16m and FSR22 (**Figure S3**). These trimeric DARPins should bind the spike protein with much higher avidity. Subsequently, using lentiviruses pseudotyped with the Wuhan-1 spike protein, we determined the neutralization activity of SR16m and SR22. Monomeric SR16m and SR22 showed weak inhibitory activity against the historical Wuhan-1 virus with 50% inhibitory concentration (IC_50_) values of 11 μM and 0.7 μM, respectively, which were enhanced 35,000- and 3,800-fold upon trimerization (**Figure 1A and B**). Despite the use of the Wuhan-1 spike protein during DARPin engineering, both FSR16m and FSR22 exhibited greater neutralization activity toward viruses pseudotyped with spike proteins from VOCs. As an example, the IC_50_ values of FSR16m and FSR22 against pseudoviruses displaying the RBD from the Omicron variant are 8.5 pM and 6.2 pM, respectively, which are 39- to 296-fold more potent than toward viruses displaying Wuhan-1 spike protein (331 pM and 1.8 nM, respectively).

**Figure 1.**
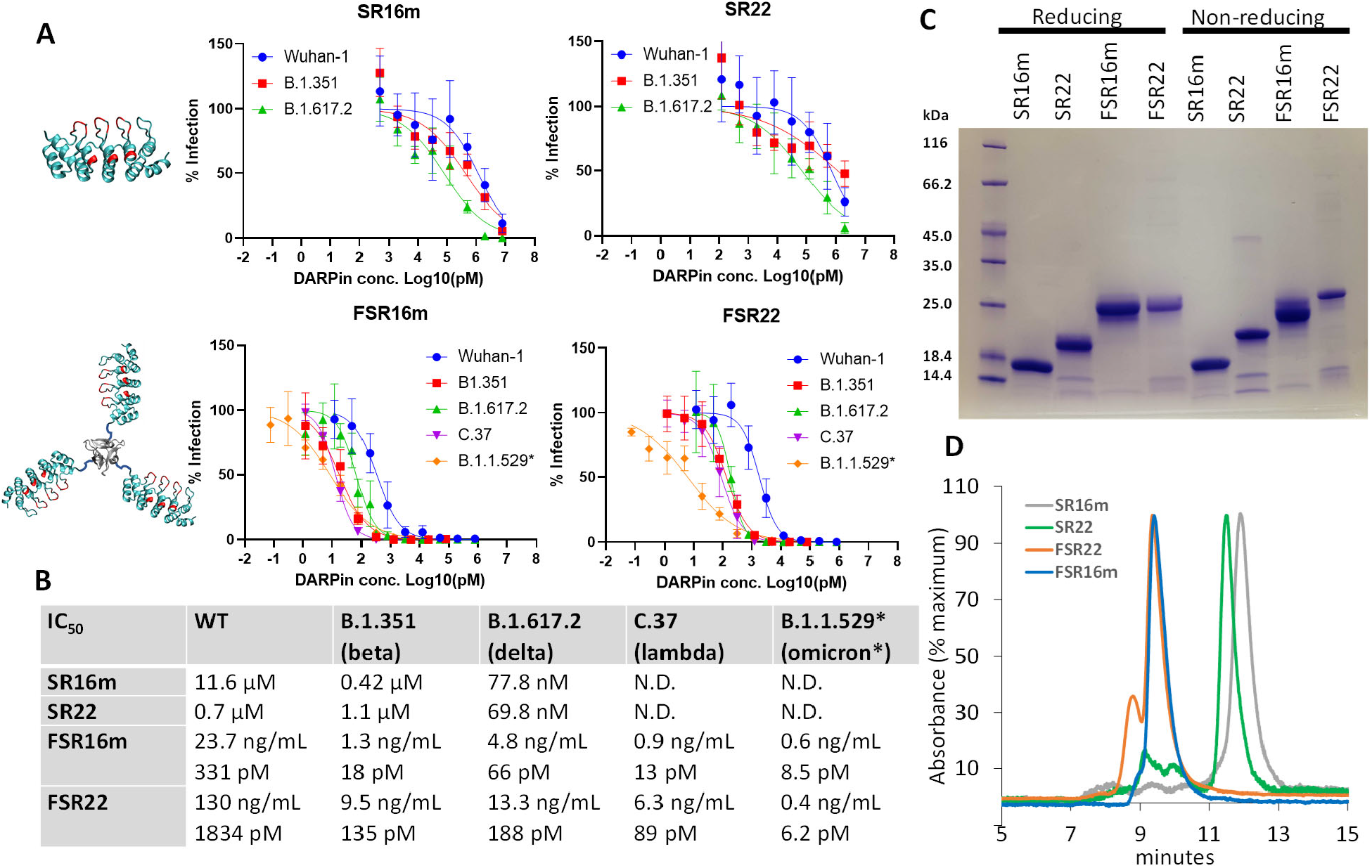
Neutralization of spike protein pseudotyped lentiviruses by DARPin molecules. (**A**) Illustration of the engineered monomeric and trimeric DARPin molecules and their neutralization of pseudotyped lentiviruses. B.1.1.529* harbors a chimeric spike protein in which the RBD region (residue 338-514) of B.1.351 was replaced with that from B.1.1.529 (BA.1). Residues in red were randomized during DARPin engineering. The error bars represent the standard deviation of at least 3 independent experiments performed in duplicate. (**B**) Summary of the neutralization IC_50_ values. (**C**) SDS-PAGE gel analysis of the DARPin molecules under non-reducing and reducing conditions. The gel image was representative of 3 experiments. (**D**) Size-exclusion chromatography of DARPin molecules under non-reducing conditions. The chromatograph was representative of 2 experiments.

All monomeric and trimeric DARPin proteins were efficiently expressed in *E*.*coli* with yields of ∼200 mg/mL and easily purified by one-step affinity chromatography. The presence of surface Cys residues (**Figure S3**) did not impact the tertiary structure of these proteins (**Figure 1C**). Monomeric DARPin SR16m migrated slightly faster than SR22 on both SDS-PAGE gel and size exclusion chromatography, while their respective trimers had nearly identical size (**Figure 1C and D**). FSR16m was homogenous in solution, whereas a small percentage of FSR22 appeared to aggregate under non-reducing conditions (**Figure 1D**).

### FSR16m and FSR22 bind diverse RBD variants with high avidity

FSR16m exhibited superior binding avidity than FSR22 against a panel of spike proteins from VOC and VOI (**Figure 2A, S4**) in a biolayer interferometry (BLI) binding assay. Due to the negligible dissociation rate, the K_Dapp_ values of FSR16m for the RBD of Wuhan-1 and 6 viral variants were below the detection limit (K_Dapp_ < 1 pM). In the case of Omicron RBD, a faster dissociation rate was observed, resulting in a K_Dapp_ of 3.65 nM. This result contrasted with our neutralization assay in which the FSR16m exhibited 39-fold improved IC_50_ values against lentiviruses displaying RBD from the Omicron variant compared to Wuhan-1 virus. The difference in the apparent binding avidity may be due to differences in the tertiary conformation of the protein in solution and on the virus. The binding study used dimeric Fc-tagged RBD. FSR22 maintained a similar binding avidity toward all tested RBDs with a slight increase in avidity to the RBD of Omicron (K_Dapp_ = 2.63 nM) compared to Wuhan-1 (K_Dapp_ = 12.3 nM). Overall, a faster dissociation rate was observed for FSR22 than FSR16m.

**Figure 2.**
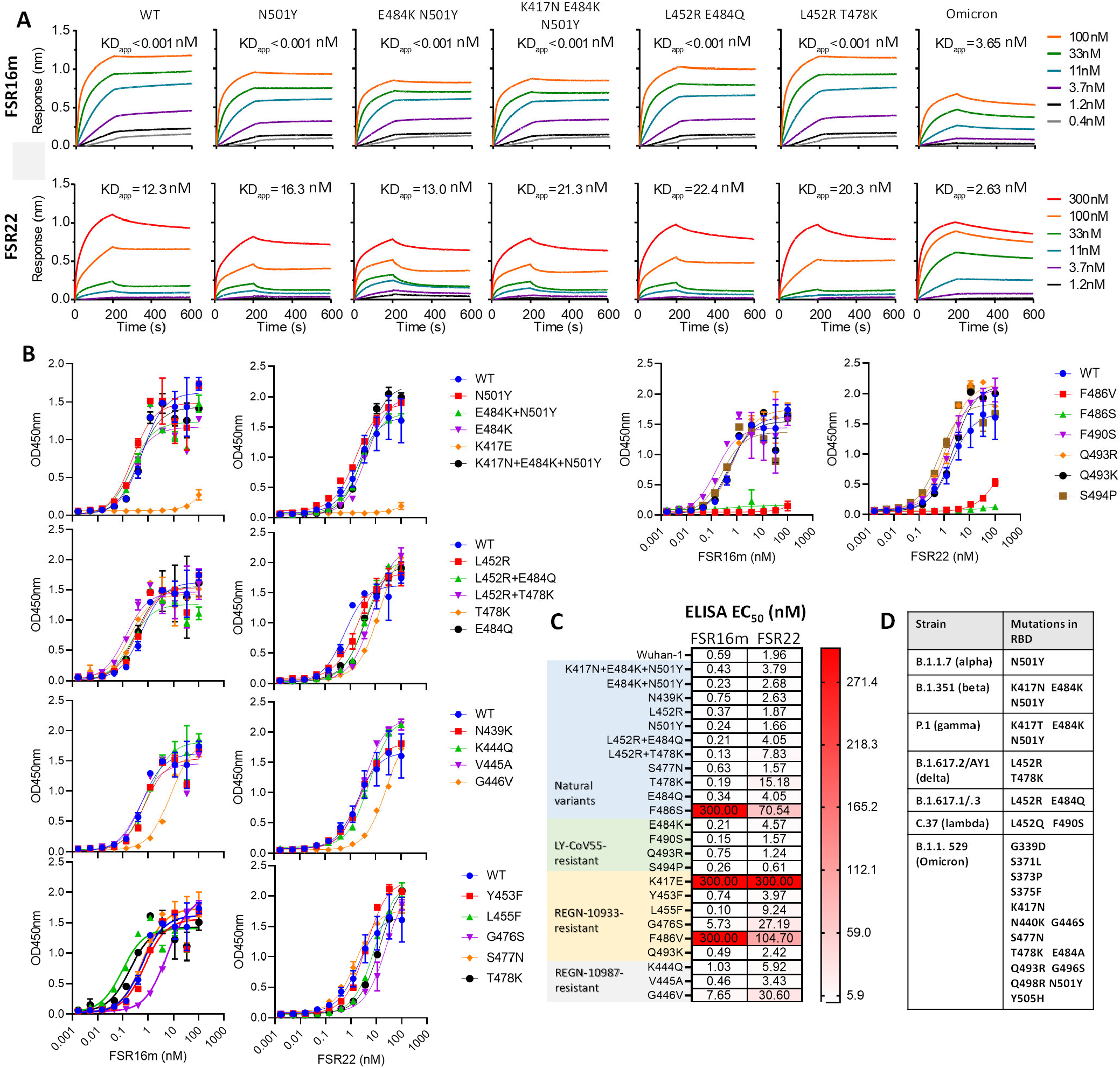
FSR16m and FSR22 maintain high avidity toward a panel of SARS-CoV2-variants. (**A**) Binding kinetics to selected mutant RBD proteins. (**B**) ELISA binding titration to a panel of 24 RBD variants. (**C**) Summary of the 50% maximum effective binding concentrations (EC_50_) to RBD proteins with indicated mutations. The EC_50_ values are the means of duplicate wells from two independent experiments. (**D**) Summary of RBD mutations.

To assess the breadth of FSR16m and FSR22 interaction with other SARS-CoV-2 VOC and VOI, we determined their binding to a panel of 24 RBD mutants by ELISA. Remarkably, FSR16m maintained nM EC_50_s toward all variants except for ones with K417E and F486V/S substitutions (**Figure 2C and D**). While the K417E mutation was predicted to escape antibody neutralization based on pseudotyped virus studies^9^, this amino acid change also results in reduced binding affinity of spike for hACE2 (17.8% of Wuhan-1 control)^10^ due to the loss of a critical salt bridge interaction between the positively-charged K417 in the spike protein and the negatively-charged Asp in hACE2. K417N/T substitutions are present in B.1.351 (Beta) and P.1 (Gamma) variants, respectively (**Figure 2D**). The affinity of FSR16m for the B.1.351 RBD (K417N+E484K+N501Y, EC_50_ 0.43 nM) is similar to Wuhan-1 RBD (EC_50_ of 0.59 nM) but slightly weaker than that to a variant with only E484K+N501Y (EC_50_ of 0.23 nM), indicating that the K417N mutation slightly weakens the interaction between FSR16m and the spike protein. Similarly, the mutations F486V and F486S also exhibit decreased hACE2 binding affinity (37% and 57%, respectively, of Wuhan- 1 control^10^). Consequently, both K417E and F486V/S mutations might adversely affect viral fitness, precluding their domination in humans. Mutations G476S and G446V enable resistance to neutralization by antibodies REGN-10933 and REGN-10987, respectively^11^. Nonetheless, and despite a greater than 10-fold increase in EC_50_ values, FSR16m still maintained nM binding avidity to spike proteins with both of these mutations (**Figure 2C**). However, no detectable interaction was observed between FSR16m and the RBD proteins of sarbecoviruses of clade 1a (SARS-CoV-1, RaTG13, WIV1), clade 2 (BtkY72) and clade 3 (Rs4081) using the same ELISA binding assay (**Figure S5**), indicating that the epitope of FSR16m is not conserved among other sarbecoviruses.

FSR22 retained high-avidity binding to most of the tested variants with EC_50_ < 10 nM, although its binding strength was generally lower than FSR16m. Similar to FSR16m, FSR22 also exhibits lower avidity to RBDs harboring mutations K417E, F486V/S, G446V, or G476S. Beyond these, FSR22 had reduced avidity to RBDs containing T478K substitution, pointing to its engagement of a slightly different binding interface than FSR16m. The ability of FSR16m and FSR22 to bind diverse RBD variants with high avidity may stem from our engineering approach, which aimed to identify binders that mimic hACE2 binding, a step that the virus is obligated to maintain as high-affinity binding for efficient infection.

### *In vivo* protection by FSR16m

We next tested the capacity of FSR16m and FSR22 to neutralize authentic SARS-CoV-2 in Vero-hACE2-TMPRSS2 cells^12^. Both FSR16m and FSR22 DARPins efficiently inhibited infection of several authentic SARS-CoV-2 strains with FSR16m exhibiting 10-fold greater potency (**Figure 3A and B**). The IC_50_ values of FSR16m against the authentic B.1.351, B.1.617.2, and B.1.617.2.AY1 viruses were 3.4, 2.2, and 3.2 ng/mL, respectively, values comparable to that of the Regen-COV antibody cocktail^13^. While both DARPins neutralized authentic Omicron strains, FSR16m was more potent. The IC_50_ values of FSR16m and FSR22 against BA.1, BA.1.1 and BA.2 strains were 44.7 and 169.2 ng/mL, 7.4 and 41.2 ng/mL, and 33.3 and 216.2 ng/mL, respectively (**Figure 3A**).

**Figure 3.**
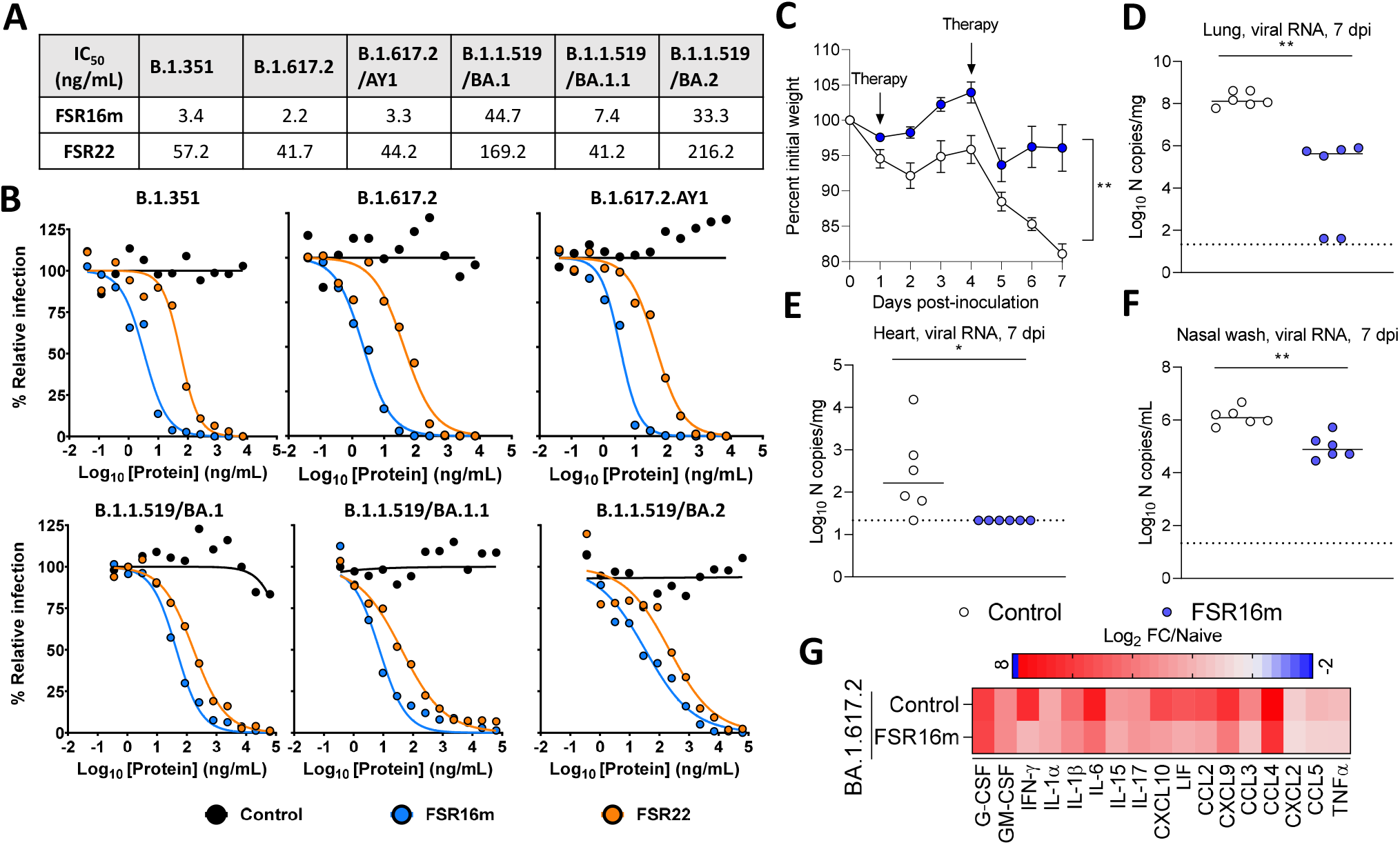
Neutralization of authentic SARS-CoV-2 viruses. (**A, B**) FSR16m and FSR22 potently neutralized authentic SARS-CoV-2 variants. DARPins were incubated with 10^2^ focus-forming units (FFU) of SARS-CoV-2 for 1 h followed by addition to Vero-hACE2-TMPRSS2 cells. One representative experiment with the mean of 2 technical replicates is shown. (**C-F**) Intranasally administered FSR16m protected mice against infection by SARS-CoV-2 B.1.617.2. (**C**) Weight change following infection with 10^3^ FFU of SARS-CoV-2 B.1.617.2 via intranasal administration. (**D-F**) Viral RNA levels at 7 dpi in the lung, heart and nasal wash (n = 6, 2 experiments). Mann-Whitney test, *p<0.05, **p<0.01. (**G**) Heat map of cytokine and chemokine protein expression levels in lung homogenates from the indicated groups. Data are presented as log_2_-transformed fold-change over naive mice. Blue, reduction; red, increase. Results are from two experiments.

Given the potency of FSR16m, we chose to investigate the *in vivo* efficacy of this DARPin in mice. Eight-week-old female heterozygous K18-hACE2 C57BL/6J^14^ mice were administered 10^3^ focus-forming units (FFU) of SARS-CoV-2 B.1.617.2 on day 0. This dose of SARS-CoV-2 was determined to cause severe lung infection and inflammation in previous studies^15-17^. On days 1 and 4 post-infection, FSR16m (50 μg/mouse in PBS) was administered via an intranasal route. FSR16m-treated mice had less weight loss and 10- to 100-fold lower levels of viral RNA in the lung, heart and nasal wash than mice treated with PBS (**Figure 3C-F**). Mice treated with FSR16m also showed lower levels of several pro-inflammatory cytokines compared to the control-treated, infected mice (**Figure 3G, S6**). Thus, FSR16m showed post-exposure therapeutic efficacy against SARS-CoV-02 as an intranasally administered protein.

### Cryo-EM structures of FSR22 and FSR16m in complex with SARS-CoV-2 spike revealed that the DARPin molecules recognize an essential component of the ACE2-binding surface

To understand the pan-neutralizing activity of FSR22 and FSR16m against SAR-CoV-2 variants, we determined their cryo-EM structures in complex with SARS-CoV-2 S6P spike proteins^18^. *Ab-initio* reconstructions of cryo-EM images of the complexes followed by heterogeneous refinements revealed trimeric SARS-CoV-2 spikes with a variety of RBD conformations, ranging from all RBD-down to one, two, or three RBD-up conformations – with SR22 or SR16m binding to the RBD-up forms in all cases (**Figure S7, S8, S9A and B**). As FSR22 and FSR16m, the trimeric forms of three SR22/SR16m linked by a foldon, were the most inhibitory, we focused on refining the three RBD-up conformations of SARS-CoV-2 spikes with the FSR22/16m bound to the tip of the RBD in its open conformation (**Figure 4A and B, and S9A and B**). Overall, the trimeric spike complexes were well-defined except for the tip of the RBD and the FSR22/FSR16m due to conformational heterogeneity. As a result, we could not delineate the FSR22/FSR16m-RBD interface in atomic detail, even after extensive heterogeneous refinement followed by local refinement (**Figure S7 and S8**). However, the overall fold of the DARPins and the tip of the RBDs fit well, revealing FSR22 and FSR16m bound to the “tip” of the RBD in their “RBD-up” conformation (**Figure 4A and B**). We determined the FSR22 structure in complex with SARS-CoV-2 spike at higher resolution than the FSR16m structure (**Figure S7 and S8**). Furthermore, with the overall fold of the two DARPins and their binding mode to SARS-CoV-2 spike being nearly identical, we used the FSR22 structure to model the FSR16m residues that differ from FSR22 (**Figure 4C and D**). Despite the limited resolution, the FSR22-bound spike suggested that residues Pro45, Ser46, Val78, Trp79, Arg81, Phe89, Asn112, Tyr114, Leu119, Arg123 and Phe145 of SR22 (**Figure S9C**) interacted with spike residues Leu455, Phe456, Glu484, Phe486, Asn487, Tyr489, and Phe490 on RBD (**Figure 4C and E**). Similarly, residues Leu45, Phe46, Ala78, Phe79, Arg81, Leu89, Tyr112, Val114, Leu119, Leu123 and Phe145 on SR16m (**Figure S9D and E**) potentially contacted spike residues Lys417, Tyr421, Leu455, Phe456, Gln474, Glu484, Gly485, Phe486, Asn487, Tyr489, and Phe490 on RBD (**Figure 4D and E**). These data explain the reduced binding of FSR16m and FSR22 for spike proteins with K417E and F486V/S mutations (**Figure 2C and D**). Of note, the FSR22/FSR16m contacting surface encompassing residues Leu455, Phe456, Ala475, Glu484, Gly485, Phe486, Asn487, Tyr489 and Phe490 overlaps closely with the hACE2-binding surface^19,20^ (**Figure 4E**). Moreover, residues Tyr421, Phe456, Phe486, Asn487 and Tyr489, which make up roughly two-thirds of the entire DARPin-interactive surface, did not overlap with residues that changed in variants of concern (VOC), including N501Y in B.1.1.7 (Alpha), K417N, E484K, and N501Y in B.1.351 (Beta), L452R and E484Q in B.1.617.2 (Delta), and K417N, S477N, Q498R, and N501Y in B.1.1.529 (Omicron) (**Figure 4E-G**), suggesting that the surface recognized by FSR22/16m is particularly important for ACE2 binding, and changes in these residues may compromise viral infectivity and fitness^20^. Comparison of the two DARPin-RBD complexes and the ACE2-RBD structure revealed a steric clash between a monomer of SR22/SR16m and ACE2 bound to RBD, as the DARPins and ACE2 recognize overlapping epitopes (**Figure 4F and G**). However, SR22 and SR16m only buried approximately 470 Å^2^ of surface area on RBD, whereas ACE2 binding buried approximately 970 Å^2^ of the surface on RBD, explaining the structural basis for the relatively poor neutralizing ability of SR22/SR16m (monomers) as they could be outcompeted by ACE2 for RBD binding (**Figure 4F and G**). Conversely, FSR22 and FSR16m, composed of three SR22 and SR16m, respectively, can overcome the lower affinity of monomeric DARPins to RBD by binding three RBDs in their open conformation (**Figure 4A and B**). Collectively, the cryo-EM structures of FSR22 and FSR16m-bound SARS-CoV-2 spike revealed that FSR22 and FSR16m potently neutralize SARS-CoV-2 including VOC by targeting a minimal but essential subset of ACE2-binding residues and taking advantage of avidity gained by trimerization (**Figure S6F**).

**Figure 4.**
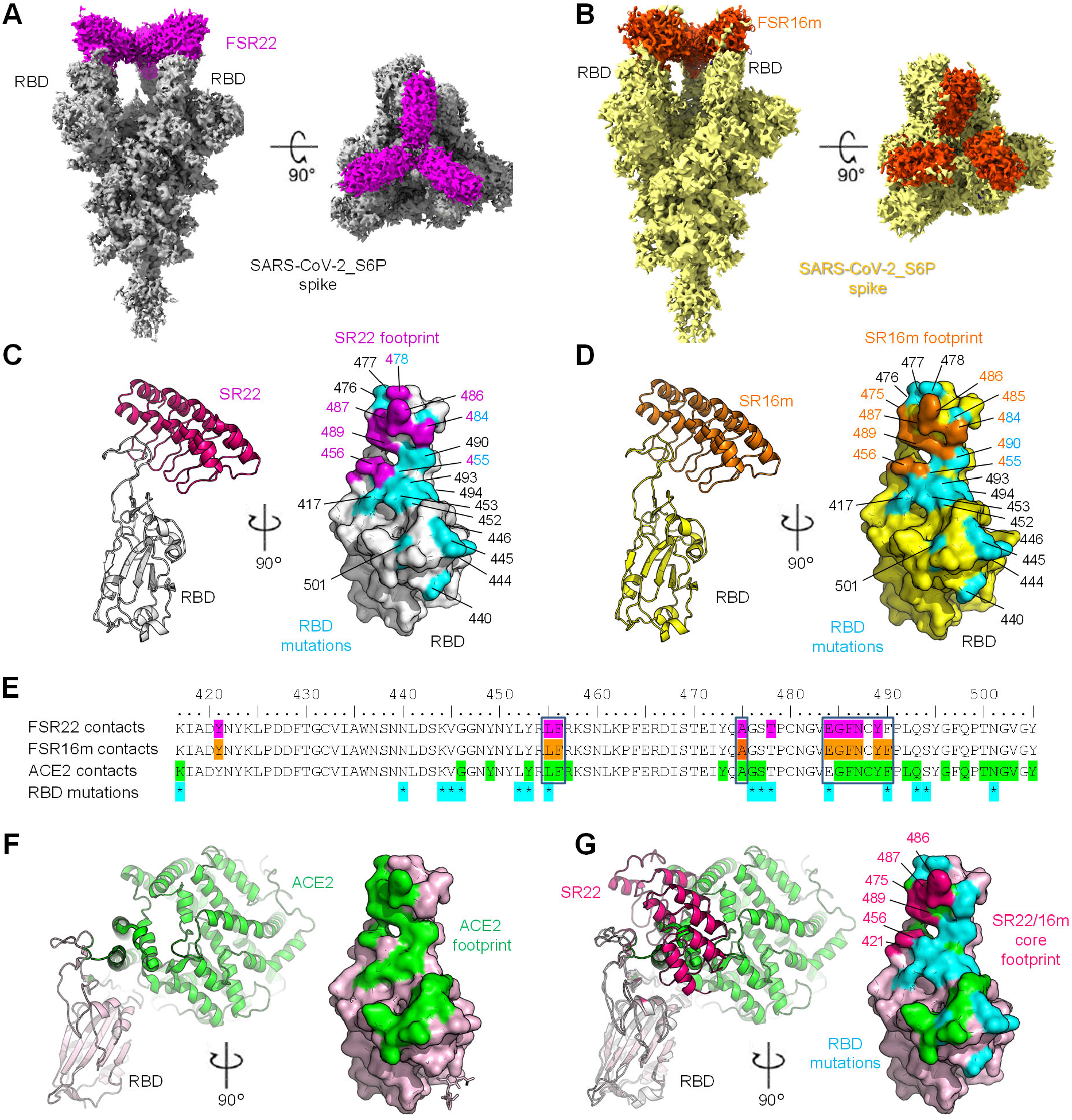
Cryo-EM structures of FSR22 and FSR16m. (**A**) Cryo-EM map of FSR22 (magenta)-bound SARS-CoV-2_S6P spike in its RBD-open conformation (gray) (**B**) Cryo-EM map of FSR16m (orange)-bound SARS-CoV-2_S6P spike (yellow). (**C**) SR22-bound RBD in cartoon representation (left). The footprint of SR22 on the surface of RBD (gray) is shown in magenta (right). RBD mutations were highlighted in cyan (right). (**D**) SR16m-bound RBD in cartoon representation (left). The footprint of SR16m on the surface of RBD (yellow) is shown in orange (right). (**E**) Contact residues on RBD by the FSR22-, FSR16m-, and the ACE2-binding are highlighted in magenta, orange, and green, respectively. RBD residues mutated in VOC are denoted with asterisks (*). (**F**) ACE2-bound RBD in cartoon representation (PDB ID:7C8D) (left). The ACE2 footprint on RBD is highlighted in green (right) (**G**) Overlay of SR22-bound RBD and ACE2-bound RBD structure (left). Overlay of footprints of ACE2, SR22/SR16m core, and RBD mutation found in variants of concern (right).

## DISCUSSION

By using phage panning coupled with functional screening, we report the engineering of DARPins with potent and broad neutralizing activity against SARS-CoV-2. Monomeric DARPin SR16m and SR22 exhibited weak viral neutralization potency, but this was enhanced 35,000- and 3,800-fold, respectively, upon trimerization by fusion with a T4 foldon. The best DARPin, FSR16m, neutralized authentic SARS-CoV-2 B.1.617.2 (Delta) with an IC_50_ of 2.2 ng/mL, similar to currently used therapeutic antibodies^21^. Although the Wuhan-1 spike protein was used for engineering, FSR22 and FSR16m exhibited 39-296-fold increased neutralization potency against viral variants, and potently neutralized pseudotyped viruses displaying chimeric spike protein with B.1.1.529 (Omicron) RBD with IC_50_ of 8.5 pM and 6.2 pM, respectively.

Our cryo-EM structural analyses revealed that both FSR22 and FSR16m recognize a distinct region of the ACE2-binding surface on RBD, particularly residues 421, 456 and 485-489. Shang et al. demonstrated that mutations of residues 481-489 reduced ACE2 binding substantially^20^, and Starr et al. reported that mutations in residues 421, 475 and 487 reduced RBD expression and possibly viral fitness, whereas mutations in residues 421, 456, 475, 486, 487 and 489 reduced ACE2 binding affinity^10^. The precise targeting of essential ACE2 interacting residues on RBD by FSR22 and FSR16m may explain their pan-neutralization of naturally derived SARS-CoV-2 variants, and our two DARPins may effectively prevent infection of variants that require hACE2 for entry.

The ability of these DARPins to exhibit increased neutralization potency toward SARS-CoV-2 variants contrasts with many human-derived anti-SARS-CoV-2 mAbs, which lose neutralization potency^3^. Multiple reasons may account for this phenomenon. (a) Naturally derived viral variants are selected in part by their ability to evade the neutralization by circulating human antibodies, including those used as human therapy^3^. This contrasts with *in vitro* engineered DARPins whose potency appears not affected by the virus evolution history. (b) The dimeric structure of an antibody limits a maximum of two spike proteins within a spike trimer to be simultaneously engaged by a single antibody, necessitating a high affinity between the antibody and the spike protein and a close match of the binding interface. Even small changes at or near the binding interface could disrupt antibody-spike protein interactions and reduce the antibody binding and neutralization potency. In contrast, due to its much smaller size, a trimer DARPin can easily engage all three monomers in a spike trimer concurrently. The strong avidity effect from the trimer interaction can compensate for the poor binding affinity of the monomer and enable the trimers to tolerate mutations at or near the binding interface without compromising potency. (c) DARPins SR16 and SR22 were selected based on their ability to compete with hACE2 for binding to the spike protein rather than their ability to inhibit viral infection. An effective way to inhibit ACE2 binding to spike is to occupy the interface for hACE2 binding, and indeed both DARPins engage key residues within the hACE2 binding interface. As emerging natural VOC tend to exhibit higher infectivity and ACE2 binding affinity^22-24^, these variants may become more susceptible to binding and neutralization by our DARPin molecules.

The upper airway epithelium is a first site of infection by SARS-CoV-2 due in part to its robust expression of hACE2^25^. Infected upper airway epithelium cells release progeny viruses, which lead to infection of other organs including the lung. Antibody therapeutics have been an effective weapon for treating COVID-19, with several virus-neutralizing IgG antibodies approved for emergency use by the FDA^26-28^. However, antibodies have several limitations as therapeutics for treating SARS-CoV-2: (a) antibody production requires sophisticated mammalian cell culture and is expensive with limited global manufacturing capacity; (b) IgG antibody requires subcutaneous or intravenous routes of administration that may be inconvenient or impractical for patients and care providers; (c) intravenously and subcutaneously administered antibodies have poor access to mucosal compartments with an estimated 50- to 100-fold lower antibody levels than in blood^29-31^, necessitating a high therapeutic dose; and (d) many current COVID-19 antibody therapeutics have a narrow neutralization spectrum and require cocktails to maintain efficacy toward newly emerged SARS-CoV-2 variants. The most recent Omicron variant was resistant to bamlanivimab, etesevimab and REGEN-COV (casirivimab and imdevimab)^32-35^.

Through the use of the stringent K18-hACE2 mouse model of SARS-CoV-2 pathogenesis, we showed that intranasal administration of FSR16m on days 1 and 4 post-infection reduced viral burden and weight loss. As several other studies have demonstrated^15,36-38^, overexpression of hACE2 in these mice results in severe lung inflammation and disease upon infection with SARS-CoV-2. Therefore, the level of protection we observed by intranasal administration of FSR16m is significant. However, the current format of intranasal delivery likely is not directly translatable to humans. To increase the drug exposure, nasally administered FSR16m may require formulation and aerosolization using a nasal spray or a handheld nebulizer. Several protein-based antivirals are currently under development against COVID-19^31,39-42^. However, unlike FSR16m, which potently neutralized all tested VOCs, many of these have narrower spectrum of neutralization^32-35^. FSR16m is efficiently expressed in *E. coli*. (>200 mg per liter of shaker flask culture, on par with that reported previously^43^), enabling cost-effective production in large-scale. In addition, FSR16m exhibits remarkable stability with <10-fold loss in activity after storage at room temperature for 6 weeks (**Figure S10**), consistent with previously reported DARPin storage stability^44^ and making the molecule a promising therapeutic candidate.

Previously, two highly potent anti-SARS-CoV-2 hetero-trimeric DARPin molecules were reported^6,7^. One of these DARPins, MM-DC (MP0420, ensovibep), binds three distinct epitopes in RBD, whereas the other, MR-DC (MP0423), targets one epitope on RBD, one on the S1 N-terminal domain, and another on the S2 domain of the spike protein. Proline-threonine-rich polypeptide linkers with designed lengths were used to connect the different DARPin molecules. Both MM-DC and MR-DC exhibited picomolar IC_50_ values against spike protein pseudotyped vesicular stomatitis virus (VSV) particles and were 10-50-fold more potent than their constituent monomer DARPins. This level of potency enhancement contrasts with both FSR16m and FSR22, whose IC_50_ values improved 35,000- and 3,800-fold, respectively, upon homo-trimerization. When administered via intraperitoneal route at 10 mg/kg, MR-DC reduced the viral load in nasal turbinates and lung by ∼10 and ∼100-fold, respectively^6^. A similar level of viral load reduction in these tissues was achieved by intranasally-administered FSR16m at 2.5 mg/kg per animal. Despite targeting three distinct regions on the spike protein, MR-DC (MP0423) lost efficacy against B.1.351, B.1.1.7, and P.1 strains^7^. MM-DC (MP0420, ensovibep) retained efficacy against all these variants and its IC_50_ toward authentic B.1.351 (7.5 ng/mL) is similar to that of FSR16m (3.4 ng/mL). MM-DC had <2-fold reductions in potency against the B.1.351 and the B.1.1.529 (Omicron) variant^45^. In contrast, the potency of FSR16m toward pseudovirus displaying RBD of B.1.351 and B.1.1.529 was increased by 18- and 38-fold, whereas the potency of FSR22 was increased by 13- and 295-fold, respectively. Thus, DARPins reported previously and those described here show differences in their recognition of VOC. Both MM-DC and MR-DC are pentameric molecules composed of two albumin binding DARPins and three spike-binding DARPins and have a half-life of 4-6 days in hamsters^6^. Intravenously administered MM-DC reduced the risk of hospitalization by 78% in a phase II clinical trial ^47^. Although FSR16m and FSR22 in their current form are expected to have a short half-life *in vivo* and are not suitable for systematic application, the same albumin binding DARPin molecules can be fused to FSR16m to extend its *in vivo* half-life and make it suitable for intravenous application.

Although DARPin molecules have low immunogenicity, the trimerization domain T4 foldon is of non-human origin and may be immunogenic in humans if administered systemically. However, since systemic absorption of intranasally-administered FSR16m is anticipated to be low due to its large size^48^, FSR16m should remain effective during the short treatment window. Nevertheless, efforts to replace T4 foldon with an equivalent trimeric protein of human origin are underway.

In summary, we report the engineering of two homotrimeric DARPin molecules, FSR16m and FSR22, with potent and broad SARS-CoV-2 neutralization activity. Both proteins are amenable to rapid and cost-effective production in *E. coli*, and intranasally administered FSR16m protected mice previously infected with SARS-CoV-2, suggesting that these proteins might be effective countermeasure candidates for COVID-19. Finally, our work suggests that homo-multimerization of monomeric proteins with moderate antiviral activity may represent an effective strategy for generating broadly neutralizing antiviral agents.

## Methods

See supplementary materials.

## Data availability

Datasets (raw data) underlying the figures have been provided as Source Data. CryoEM structures are deposited in the protein data bank with pdb codes 7TYZ and 7TZ0.

## Acknowledgments

Prof. Paul Bieniasz provided the 293T clone 22 cells for pseudoparticle neutralization assays. Prof. Nathaniel Landau provided the plasmids for Δ19 spike protein of B.1.617.2 and C.37. Dr. Aaron Benjamin assisted with Size Exclusion Chromatography experiments. The following reagent was obtained through BEI Resources, NIAID, NIH: Spike Glycoprotein (Stabilized) from SARS-Related Coronavirus 2, Wuhan-Hu-1 with C-Terminal Histidine and Twin-Strep® Tags, Recombinant from HEK293 Cells, NR-52724.

## Author contribution

Conception: VC, MSD, PDK, ZA, ZC; Methodology: VC, YDK, JG, JBC, ZK, RS, HB, FK, MSD, PDK, ZA, ZC; Investigation: VC, YDK, JG, JBC, ZK, RS, RGC, DRH, ASO, TS, HB, YT; Funding Acquisition: FK, MSD, PDK, LS, ZA, ZC.

