## Supplementary information for "Potent and pan-neutralization of SARS-CoV-2 variants of concern by DARPins"

**Supplementary material:**

**Methods**

***Selection and screening of spike protein-binding DARPin molecules***

An in-house N3C DARPin library with >$10^9$ diversity was used in the phage panning essentially as described previously[1]. Purified full length spike protein (BEI 52724) and RBD (BEI52306) were biotinylated via EZ-link-sulfo-NHS-LC-biotin (Thermo Fisher, Cat #21335) and used as target proteins. RBD was used as the target protein in Rounds 1, 2 and 4 while the full-length spike protein was used as the target in Round 3 to ensure the enrichment of DARPins that recognize RBD present on the full-length spike protein. Round 1 used the target protein (100 nM) in solution whereas Rounds 2-4 employed decreasing concentrations of the target protein immobilized on streptavidin-coated ELISA plate (100 nM, 50 nM and 20 nM). The enrichment of RBD binding DARPins was confirmed by phage ELISA against both RBD and full-length spike protein following a published protocol[1, 2] (**Figure S1**).

The enriched DARPin pool from the 4$^{th}$ Round was cloned into the pET28a vector for high-level DARPin expression as described previously[3]. The resulting DARPin contains a Myc tag at the N-terminus and a 6xHis tag at the C-terminus. After transformation, a total of 300 individual *E. coli* BL21 (DE3) clones were picked and grown in deep 96-well plates (1 mL/well) at 37 ˚C in LB and induced with IPTG (0.5 mM) The next day cell pellets were harvested, resuspended in 200 µL of PBS (1.8 mM $KH_2PO4$, 10 mM $Na_2HPO4$, 137 mM NaCl, 2.7 mM KCl, pH 7.4) supplemented with lysozyme (200 µg/mL, Amresco Cat #0663-5G)[3]. An ELISA was used to identify the target binding DARPins. Briefly, Nunc MaxiSorp plates (Fisher scientific Cat #50-712-278) were coated with 4 µg/mL neutravidin (Thermoscientific, Cat #31000) in PBS at room temperature for 2 hours. The wells were washed with PBS and then blocked with PBS supplemented with 0.5% BSA (Fisher Scientific Cat #BP9706100, PBS-B) at 4˚C overnight. The next day, after washing with PBS-T (PBS with 0.1% Tween 20), the wells were incubated with biotinylated RBD or full-length spike protein (20 nM in PBS) at room temperature for 1 hours. The wells were washed again with PBS-T prior to the addition of 100 µL of cell lysate (5-fold diluted in PBS) and incubated at room temperature for 2 h. The amount of plate bound DARPin in each well was quantified using mouse anti-myc antibody (Invitrogen, Cat #13-2500, 1:2500 diluted in PBS-B) and HRP conjugated goat anti-mouse antibody (Jackson ImmunoResearch catalog # 115-035-146, 1:1000 diluted in PBS-B) as the primary and secondary antibody, respectively, and BioFx TMB (VWR, Cat #100359-154) for color development. 20 and 23 of clones showed significant ability to bind to RBD and full-length spike protein, respectively. Sequencing revelated 11 unique clones with significant ability to bind both RBD and full-length spike protein.

To identify DARPin clones able to block spike protein and hACE2 interaction, a competitive ELISA was used[4]. Briefly, the Maxisorp plates were first coated with full-length spike protein (BEI 52308, 2 nM in PBS) at room temperature for 2 h. The wells were blocked with PBS-B at 4 ˚C overnight, washed with PBS-T, and then incubated with mixtures of hACE2-HRP (prepared in-house, 0.5 nM) and different IMAC-purified DARPins (50 mM)[3, 5] at room temperature for 1 h. The amount of hACE2-HRP in each well was quantified using BioFx TMB.

For preparation of hACE2-HRP, hACE2 (Raybiotech, Cat #230-30165, 1.76 mg/mL) was first biotinylated as described above and then incubated with an equal molar amount of streptavidin-HRP (JIR Cat# 016-030-084) in PBS at room temperature for 20 min and stored at -20 ˚C in 50% glycerol until use.

**Plasmids**

Plasmids encoding the wild type Δ19 spike protein was obtained from Addgene (Cat # 145780). The plasmids for the Δ19 spike protein of B.1.617.2 and C.37 were generously provided by Prof. Nathaniel Landau[6]. DNA fragment encoding the Δ19 spike protein of strain B. 1.351 and the RBD (residues 339-501) of B.1.1.529 was synthesized by Gene Universal and inserted into the pCG1 plasmid[7]. Prof. Paul Bieniasz provided the 293T cell clone 22 (293T.c22) with high expression efficiency of human ACE2 and the lentiviral reporter plasmid pHIV-1NL4-3-ΔEnv-NanoLuc[8]. The plasmid encoding the chimeric

B.1.1.529 spike protein (B.1.1.529*) was constructed by replacing the RBD region (residues 338-514) in B.1.351 with that from B.1.1.529. Briefly, the gBlock fragment of B.1.1.529 RBD was digested with BsaI and BlpI, and ligated to 1) pCG1-B.1.351 backbone digested with BamHI and BlpI and 2) PCR product amplified from the same backbone with primers Spike_F and Spike_R and digested with BamHI and BsaI in a three-fragment-ligation reaction.

gBlock of B.1.1.529 RBD:

Aaaaaaggtctcacttcgatgaggtgttcaatgccaccagattcgcctctgtgtacgcctggaaccggaagcggatcagcaattgcgtggccgactactccgtgct
gtacaacctggccccttttcttcaccttcaagtgctacggcgtgtcccctaccaagctgaacgacctgtgcttcacaaacgtgtacgccgacagcttcgtgatccggg
gagatgaagtgcggcagattgcccctggacagacaggcaacatcgccgactacaactacaagctgcccgacgacttcaccggctgtgtgattgcctggaacagc
aacaagctggactccaaagtctctggcaactacaattacctgtaccggctgttccggaagtccaatctgaagcccttcgagcgggacatctccaccgagatctatc
aggccggcaataagccttgtaacggcgtggccggcttcaactgctacttcccactgagatcctactcctttagacccacatatggcgtgggccaccagccctacag
agtggtggtgctgagcttcgaa

BsaI BlpI Omicron mutations

(chronological order – G339D, S371L, S373P, S375F, K417N, N440K, G446S, S477N, T478K, E484A, Q493R, Q496S, Q498R, N501Y, Y505H)

Spike_F: cgaattc*ggatcc*gccacca (contains BamHI)

Spike_R: DDDDDD..D'D'D.##....'#'#.#DD..D.# (Contains BsaI)

To construct trimeric DARPin molecules, the DNA fragment encoding a codon optimized T4 foldon (pdb:1rfo) was synthesized by Gene Universal. T4 foldon was fused to the N-terminus of a DARPin molecule via a flexible (GGGGSLQ)x2 linker and cloned into the pET28a expression vector. DARPin SR16, SR22, FSR16 and FSR22 contain a 6xHis tag and a Myc tag at the N-terminus, while SR16m and FSR16m contains only a 6xHis tag at the C-terminus.

**SARS-CoV-2 spike lentiviral pseudoviruses**

Lentiviral pseudoviruses with different SARS-CoV-2 spike proteins ($CoV2_{pp}$) were produced as previously reported[8]. Briefly, plasmids encoding the Δ19 spike protein and reporter pHIV-1NL4-3-ΔEnv-NanoLuc[8] (1:3 molar ratio, 10 µg total) were mixed with 500 µL of serum-free DMEM medium and 44 μL of PEI (1 mg/mL, Polysciences transporter 5, Cat# 26008-5) and used to transfect 5 x $10^6$ 293T cells seeded the night before. 24 hours post transfection, the medium was replaced with fresh DMEM supplemented with 10% FBS and 48 hours post transfection the viral supernatant was harvested, aliquoted and stored at -80 ˚C until use.

To determine the neutralization efficiency, serially diluted DARPin molecules were incubated with $CoV2_{pp}$ (final 500-fold diluted) at 37˚C for 30 min before being added to 293T.c22 cells seeded the night before at $10^4$ cells/well in 96 well plates. The plates were incubated at 37˚C/5% $CO_2$ for 48 h, and the NanoLuc signal from each well was quantified using the Nano-Glo Luciferase Assay kit (Promega Cat # N1120).

**DARPin protein production and characterization**

All DARPin molecules were expressed in *E. coli* Bl21 (DE3) cells in LB medium supplemented with 50 μg/mL of kanamycin. Protein expression was induced with IPTG (0.5 mM) when the culture reached OD600~0.5. The protein expression was continued at 37˚C for 5 h, and the cells were harvested by centrifugation. The proteins were purified using gravity Ni-NTA agarose columns. Protein purity was determined using 12% SDS-PAGE gels.

For *in vivo* studies, the IMAC purified FSR16m was sterilized by filtration through a 0.22 μm filter, concentrated and buffer exchanged into PBS via ultrafiltration (Amicon column MWCO 10 KDa, Cat # UFC801024) before endotoxin removal using High Capacity Endotoxin Removal Spin Columns (Pierce Cat # 88274). Endotoxin level in the protein sample (1.5 mg/mL) was quantified to be <30 U/mL using Pierce Chromogenic Endotoxin Quant Kit (ThermoFisher Cat# A39552).

For size exclusion chromatography studies, DARPin samples (0.9 mg/mL x 0.25 mL) were loaded onto an Enrich SEC 70 x 300 Column equilibrated with PBS (GE AKTApure). Vitamin B12 (Sigma V2876-100MG) was used at a final concentration of 1 mg/ml as an internal control.

**Spike RBD and DARPin avidity measurement**

The human IgG1 Fc-tagged RBD proteins were made in-house as previously described[9, 10]. The avidity measurement was performed on the ForteBio Octet RED 96 system (Sartorius, Goettingen, Germany). Briefly, the RBD proteins (20 µg/ml) were captured onto protein A biosensors for 300 s. The loaded biosensors were then dipped into the kinetics buffer for 10 s for adjustment of baselines. Subsequently, the biosensors were dipped into serially diluted (from 0.13 to 300 nM) DARPin proteins for 200 s to record association kinetics and then dipped into kinetics buffer for 400 s to record dissociation kinetics. Kinetic buffer without DARPin was used to correct the background. The Octet Data Acquisition 9.0 software was used to collect affinity data. For fitting of $K_D$ values, Octet Data Analysis software V11.1 was used to fit the curve by a 1:1 binding model using the global fitting method.

**ELISA binding assay**

ELISA plates were coated with recombinant DARPin protein (2 µg/ml) at 4 °C overnight and blocked with 5% skim milk at 37 °C for 2 h. One hundred µL of serially diluted human IgG1 Fc-tagged RBD proteins in 1% skim milk was added to each well, and the plates were incubated at room temperature for 3 h before the addition of 100 µL/well of HRP-conjugated goat anti-human IgG antibody (Jackson ImmunoResearch, 109-035-088, diluted 1:5,000) and the plates were incubated at room temperature for another hour. The plates were washed 3 to 5 times with PBST (0.05% Tween-20) between incubation steps. TMB (3,3′,5,5′-tetramethylbenzidine) substrate was added at 100 µl per well for colour development. The reaction was stopped by adding 50 µl per well 2M $H_2SO_4$. The OD450 nm was read by a SpectraMax microplate reader and analysed with GraphPad Prism 8.

**Cells and authentic viruses**

Vero-TMPRSS2[11] and Vero-hACE2-TMPRSS2[12] (a gift of A. Creanga and B. Graham, NIH) were cultured at 37°C in Dulbecco's Modified Eagle medium (DMEM) supplemented with 10% fetal bovine serum (FBS), 10 mM HEPES pH 7.3, 1 mM sodium pyruvate, 1× non-essential amino acids, 100 U/ml of penicillin–streptomycin, and 5 ug/mL of puromycin. The B.1.617.2, B.1.617.2.AY1, B.1.351, BA.1, BA.1.1, and BA.2 SARS-CoV-2 strains were obtained from infected individuals and have been described previously[1314]. Infectious stocks were propagated by inoculating Vero-hACE2-TMPRSS2 cells. Supernatant was collected, aliquoted, and stored at -80$^{o}$C. All work with infectious SARS-CoV-2 was performed in Institutional Biosafety Committee-approved BSL3 and A-BSL3 facilities at Washington University School of Medicine using positive pressure air respirators and protective equipment. All virus stocks were deep-sequenced after RNA extraction to confirm the presence of the anticipated substitutions.

**Mouse experiments**

Animal studies were carried out in accordance with the recommendations in the Guide for the Care and Use of Laboratory Animals of the National Institutes of Health. The protocols were approved by the Institutional Animal Care and Use Committee at the Washington University School of Medicine (assurance number A3381–01). Virus inoculations were performed under anesthesia that was induced and maintained with ketamine hydrochloride and xylazine, and all efforts were made to minimize animal suffering.

Heterozygous K18-hACE C57BL/6J mice (strain: 2B6.Cg-Tg(K18-ACE2)2Prlmn/J) were obtained from The Jackson Laboratory. Animals were housed in groups and fed standard chow diets. 8-week-old female mice were administered $10^3$ FFU of SARS-CoV-2 via intranasal administration.

**Measurement of viral burden**

Tissues were weighed and homogenized with zirconia beads in a MagNA Lyser instrument (Roche Life Science) in 1,000 μL of DMEM media supplemented with 2% heat-inactivated FBS. Tissue homogenates were clarified by centrifugation at 10,000 rpm for 5 min and stored at −80°C. RNA was extracted using the MagMax mirVana Total RNA isolation kit (Thermo Scientific) on a Kingfisher Flex extraction robot (Thermo Scientific). RNA was reverse transcribed and amplified using the TaqMan RNA-to-CT 1-Step Kit (ThermoFisher). Reverse transcription was carried out at 48°C for 15 min followed by 2 min at 95°C. Amplification was accomplished over 50 cycles as follows: 95°C for 15 s and 60°C for 1 min. Copies of SARS-CoV-2 *N* gene RNA in samples were determined using a previously published assay [15, 16]. Briefly, a TaqMan assay was designed to target a highly conserved region of the N gene (Forward primer: ATGCTGCAATCGTGCTACAA; Reverse primer: GACTGCCGCCTCTGCTC; Probe: /56-FAM/TCAAGGAAC/ZEN/AACATTGCCAA/3IABkFQ/). This region was included in an RNA standard to allow for copy number determination down to 10 copies per reaction. The reaction mixture contained final concentrations of primers and probe of 500 and 100 nM, respectively.

**Cytokine and Chemokine protein measurements.** Lung homogenates were incubated with Triton-X-100 (1% final concentration) for 1 h at room temperature to inactivate SARS-CoV-2. Homogenates were analyzed for cytokines and chemokines by Eve Technologies Corporation (Calgary, AB, Canada) using their Mouse Cytokine Array/Chemokine Array 31-Plex (MD31) platform.

**Authentic virus neutralization assay**

Serial dilutions of DARPins were incubated with $10^2$ focus-forming units (FFU) of the indicated SARS-CoV-2 strains for 1 h at 37°C. DARPin-virus complexes were added to Vero-hACE2-TMPRSS2 cell monolayers in 96-well plates and incubated at 37°C for 1 h. Subsequently, cells were overlaid with 1% (w/v) methylcellulose in MEM supplemented with 2% FBS. Plates were harvested 24 h later by removing overlays and fixed with 4% PFA in PBS for 20 min at room temperature. Plates were washed and sequentially incubated with an oligoclonal pool of SARS2-2, SARS2-11, SARS2-16, SARS2-31, SARS2-38, SARS2-57, and SARS2-71 anti-spike protein antibodies [17] and HRP-conjugated goat anti-mouse IgG in PBS supplemented with 0.1% saponin and 0.1% bovine serum albumin. SARS-CoV-2-infected cell foci were visualized using TrueBlue peroxidase substrate (KPL) and quantitated on an ImmunoSpot microanalyzer (Cellular Technologies). Data were processed using Prism software (GraphPad Prism 8.0).

#### Cryo-EM sample and grid preparation

SARS-CoV-2 S6P spike protein was produced in 293F cells and purified from cell culture supernatant using a Ni-NTA agarose column followed by Superdex S200 16/600 (GE Healthcare) size-exclusion column chromatography as described[18]. FSR16m and FSR22 were produced in *E.coli* Bl21(DE3) cells in LB medium as described above. To generate SARS-CoV-2 S6P and FSR16m (or FSR22) complexes, SARS-CoV-2 S6P and FSR16m (or FSR22) were incubated in 1:3 molar ratio and the complexes were purified by Superdex S200 GL 10/300 (GE Healthcare) using 10 mM HEPES, 7.4, 150 mM NaCl as running buffer and were confirmed by SDS-PAGE and negative stain EM. 2.3 µl of the complex at 0.5 mg/ml concentration was deposited on a C-flat grid (protochip.com). The grids were vitrified using an FEI Vitrobot Mark IV (Themo Fisher Scientific) with a wait time of 30 s, blot times of 1.5-4.5 s and blot force of 1.

#### Cryo-EM data collection and processing

The FSR22/16m: SARS-CoV-2 S6P complexes grids were imaged using a Titan Krios electron microscope equipped with a Gatan K3 Summit direct detection device. Movies were collected at 105,000x magnification over a defocus range of -1.0 µm to -2.5 µm for a 10 s with the total dose of 58.06 e–/Å2 fractionated over 50 raw frames. All data processing was done with cryoSPRACv3.3.1. [19]. Motion correction and CTF estimation in patch mode, blob particle picking, and particle extraction with the box size of 500 Å were performed followed by 2D classifications, ab initio 3D reconstruction, and multiple rounds of 3D heterogeneous refinement. C3 symmetry was applied for the final reconstruction of the FSR22/16m: SARS-CoV-2 spike complex after the initial 3D heterogeneous refinement using C1 symmetry identified a trimer with 3 RBD-up conformation bound three SR22/16m molecules. To define RBD-FSR22 interface, local refinement was performed using a soft mask covering one SR22 and one RBD molecule.

#### Model building and refinement

Coordinates from PDB ID:7BNO [20]and the initial models of SR22/16m generated by using AlphaFold [21]were used for initial fit to the reconstructed maps. Then the models were manually built using Coot [22] followed by simulated annealing and real space refinement in Phenix [23]iteratively. Geometry and map fitting were evaluated throughout the process using Molprobity [24]and EMRinger [25]. Figures were generated using PyMOL (www.pymol.org) and UCSF ChimeraX.v1.1.1[26].

**Supplementary Figures:**

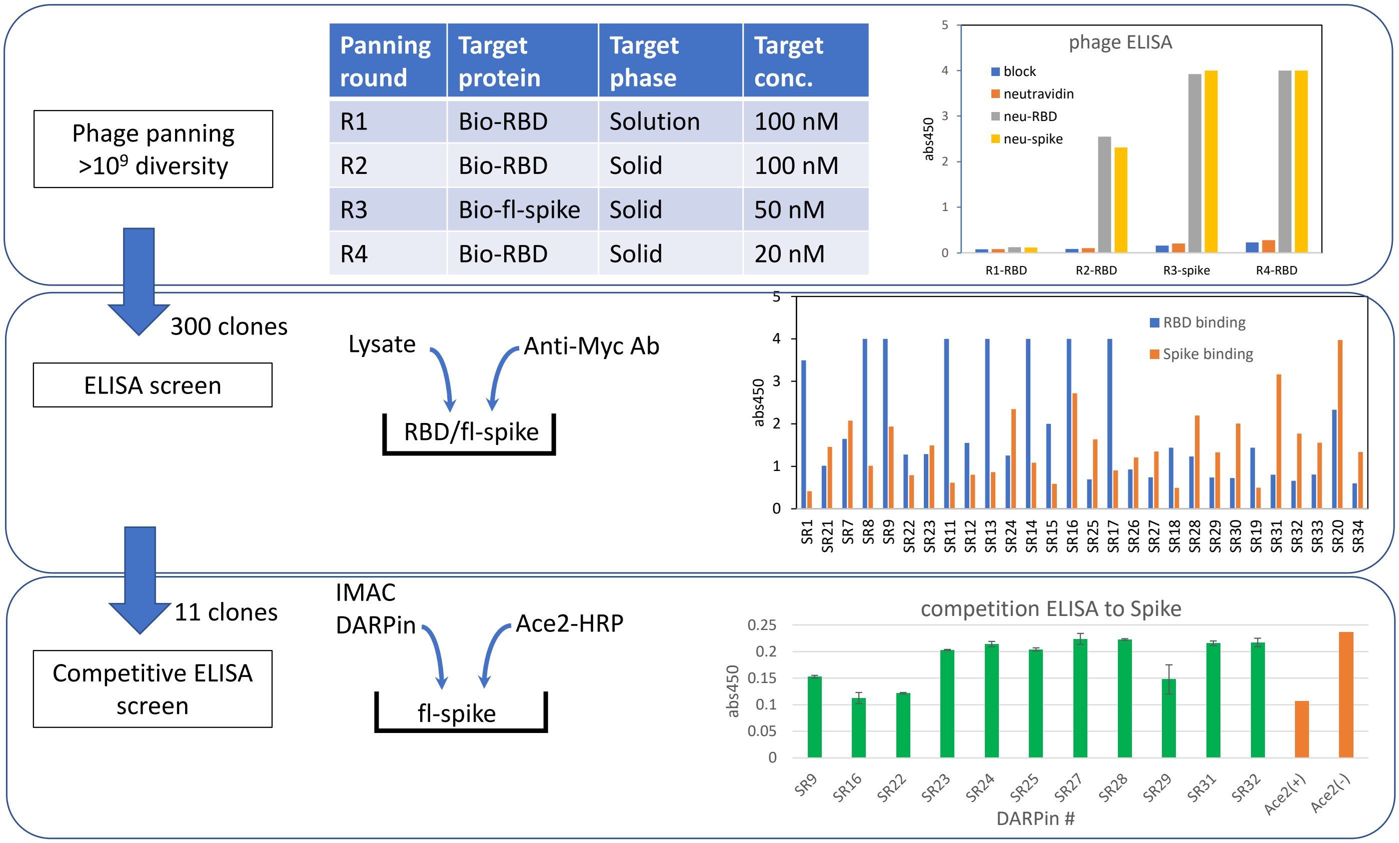


| Panning round | Target protein | Target phase | Target conc. |
|---|---|---|---|
| R1 | Bio-RBD | Solution | 100 nM |
| R2 | Bio-RBD | Solid | 100 nM |
| R3 | Bio-fl-spike | Solid | 50 nM |
| R4 | Bio-RBD | Solid | 20 nM |

**Figure S1.** Overview of the engineering procedure. Upper panel: phage panning to enrich DARPin molecules able to bind the spike protein. Middle panel: ELISA-based screen to identify DARPin variants able to bind both the RBD and the full-length spike protein. Lower panel: Competition ELISA-based screen to identify DARPin molecules able to block the binding of hACE2 to immobilized full-length spike protein.

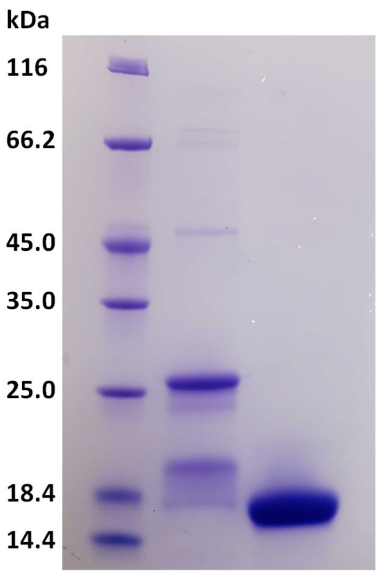


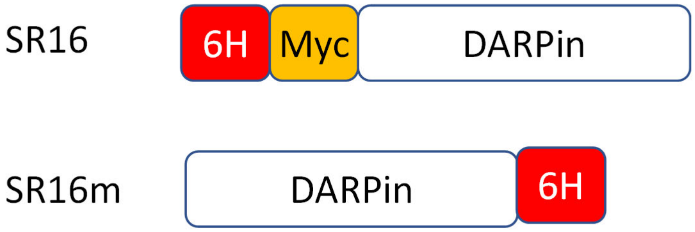


**Figure S2.** SDS-PAGE of DARPin SR16 and SR16m. The N-terminus 6xHis tag in SR16 was moved to the C-terminus to generate SR16m.

**FSR22**

MGSSHHHHHHSSGMEQKLISEEDLDGYIPEAPRDGQAYVRKDGEWVLLSTFLGGGGSLQGGGGSLQGSDLGKKLLEAARAGQ
DDEVRILMANGADVNACDPSGITPLHLAADKGHLEIVEVLLKYGADVNAMDVWGRTPLHLAAFTGHLEIVEVLLKYGADVNACD
LNGYTPLHLAAGRGHLEIVEVLLKNGAGVNAQDKFGKTAFDISIDNGNEDLAEILQSSS

**FSR16m**

MGMEQKLISEEDLDGYIPEAPRDGQAYVRKDGEWVLLSTFLGGGGSLQGGGGSLQGSDLGKKLLEAARAGQDDEVRILMANG
ADVNALDLFGVTPLHLAAERGHLEIVEVLLKNGADVNAGDAFGRTPLHLAALGGHLEIVEVLLKNGADVNACDLYGVTPLHLAAG
LGHLEIVEVLLKNGADVNAQDKFGKTAFDISIDNGNEDLAEILQSSSKLAAALEHHHHHH

**SR22**

MGSSHHHHHHSSGLVPRGSHMEQKLISEEDLGSDLGKKLLEAARAGQDDEVRILMANGADVNACDPSGITPLHLAADKGHLEIV
EVLLKYGADVNAMDVWGRTPLHLAAFTGHLEIVEVLLKYGADVNACDLNGYTPLHLAAGRGHLEIVEVLLKNGAGVNAQDKFG
KTAFDISIDNGNEDLAEILQSSS

**SR16m**

MGSSDLGKKLLEAARAGQDDEVRILMANGADVNALDLFGVTPLHLAAERGHLEIVEVLLKNGADVNAGDAFGRTPLHLAALGG
HLEIVEVLLKNGADVNACDLYGVTPLHLAAGLGHLEIVEVLLKNGADVNAQDKFGKTAFDISIDNGNEDLAEILQSSSKLAAALEH
HHHHH

**SR16**

MGSSHHHHHHSSGLVPRGSHMEQKLISEEDLGSDLGKKLLEAARAGQDDEVRILMANGADVNALDLFGVTPLHLAAERGHLEIV
EVLLKNGADVNAGDAFGRTPLHLAALGGHLEIVEVLLKNGADVNACDLYGVTPLHLAAGLGHLEIVEVLLKNGADVNAQDKFGK
TAFDISIDNGNEDLAEILQSSS

**Figure S3. Amino acid sequences of the various DARPin molecules.** T4 foldon is shaded in gray.

| FsR16m | | | | | | | |
|---|---|---|---|---|---|---|---|
| | KD (M) | KD Error | kon(1/Ms) | kon Error | kdis(1/s) | kdis Error | Full R^2 |
| WT | <1.0E-12 | 1.76E-11 | 3.85E+05 | 3.85E+03 | <1.0E-07 | NA | 0.9932 |
| N501Y | <1.0E-12 | 1.23E-11 | 4.67E+05 | 3.92E+03 | <1.0E-07 | NA | 0.9953 |
| E484K+N501Y | <1.0E-12 | 8.54E-12 | 6.39E+05 | 5.29E+03 | <1.0E-07 | NA | 0.995 |
| K417N+E484K+N501Y | <1.0E-12 | 9.97E-12 | 5.76E+05 | 4.98E+03 | <1.0E-07 | NA | 0.9948 |
| L452R+E484Q | <1.0E-12 | 1.22E-11 | 4.74E+05 | 4.03E+03 | <1.0E-07 | NA | 0.9952 |
| L452R+T478K | <1.0E-12 | 1.53E-11 | 3.99E+05 | 3.56E+03 | <1.0E-07 | NA | 0.9948 |
| Omicron | 3.65E-09 | 6.09E-11 | 1.74E+05 | 9.59E+02 | 6.36E-04 | 1.00E-05 | 0.9842 |

| FsR22 | | | | | | | |
|---|---|---|---|---|---|---|---|
| | KD (M) | KD Error | kon(1/Ms) | kon Error | kdis(1/s) | kdis Error | Full R^2 |
| WT | 1.23E-08 | 3.07E-10 | 4.61E+04 | 3.63E+02 | 5.67E-04 | 1.34E-05 | 0.9807 |
| N501Y | 1.63E-08 | 4.43E-10 | 3.77E+04 | 3.78E+02 | 6.14E-04 | 1.55E-05 | 0.9734 |
| E484K+N501Y | 1.30E-08 | 4.16E-10 | 7.96E+04 | 1.21E+03 | 1.04E-03 | 2.91E-05 | 0.8805 |
| K417N+E484K+N501Y | 2.13E-08 | 5.06E-10 | 4.56E+04 | 5.37E+02 | 9.70E-04 | 2.01E-05 | 0.9551 |
| L452R+E484Q | 2.24E-08 | 4.83E-10 | 3.83E+04 | 3.97E+02 | 8.57E-04 | 1.62E-05 | 0.9731 |
| L452R+T478K | 2.03E-08 | 4.79E-10 | 3.53E+04 | 3.61E+02 | 7.17E-04 | 1.53E-05 | 0.9773 |
| Omicron | 2.63E-09 | 4.01E-11 | 1.64E+05 | 5.80E+02 | 4.31E-04 | 6.39E-06 | 0.9938 |

**Figure S4.** The kinetic constants $K_{on}$ and $K_{dis}$, and the $R^2$, which is the coefficient of determination and an estimate of the goodness of the curve fit, for the BLI measurements shown in Figure 2A.

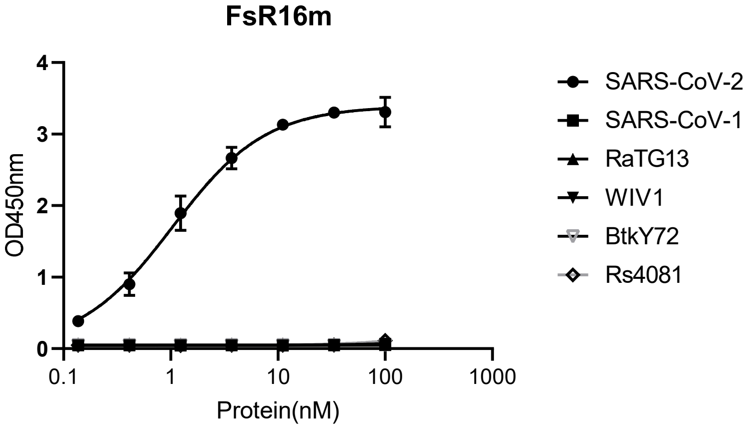


**Figure S5.** ELISA binding titration of FSR16m to the RBD proteins of five sarbecoviruses.

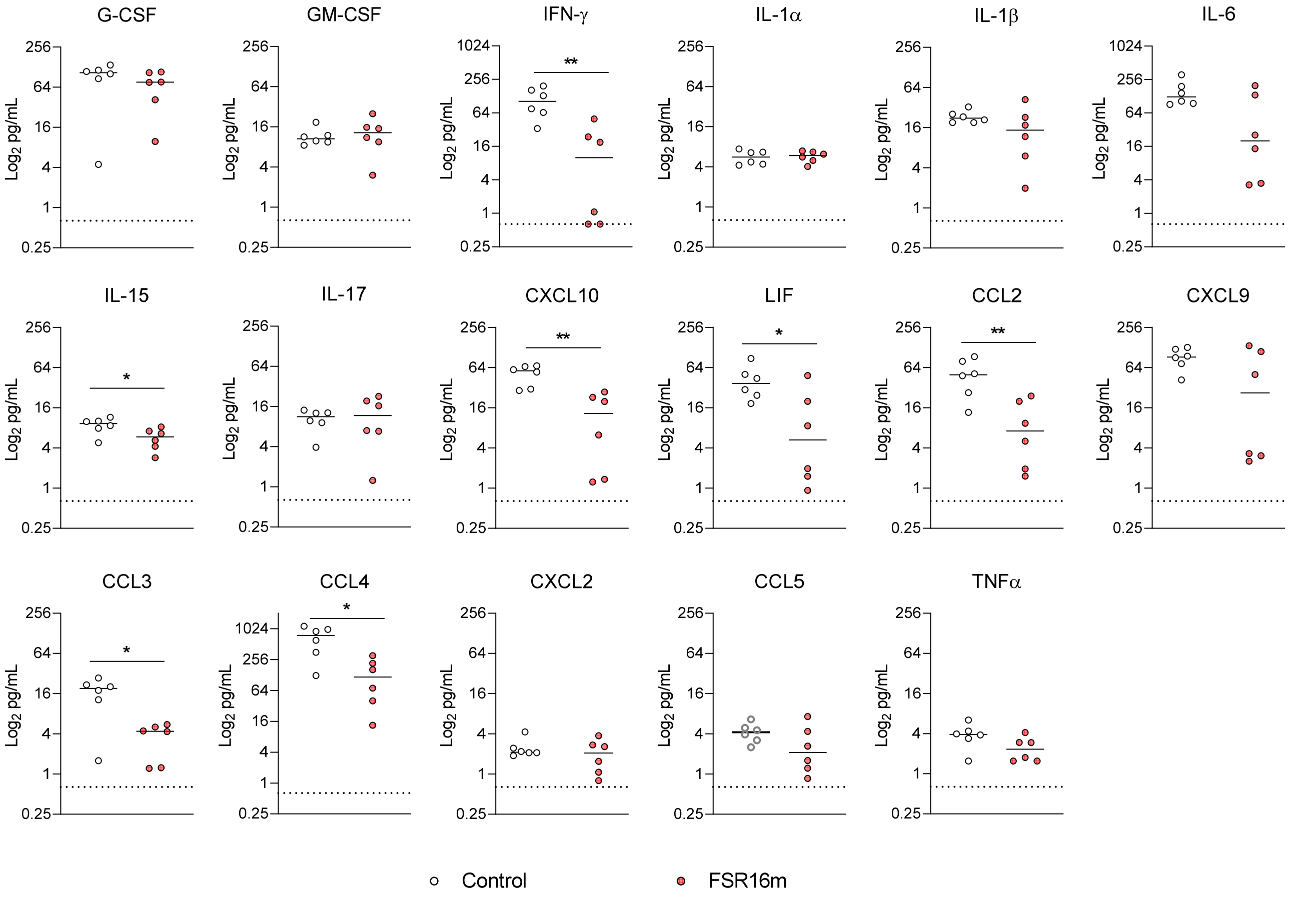


**Figure S6. Cytokine and chemokine protein concentrations in the lungs of B.1.617.2-infected mice**. Individual graphs of cytokine and chemokine protein levels in the lungs of control or FSR16m-treated K18-hACE2 mice at 7 dpi (line indicates median; n = 3 naive, n = 6 for all other groups (Mann-Whitney test between the control and FSR16m groups: *, $P < 0.05$, ** $P < 0.01$).

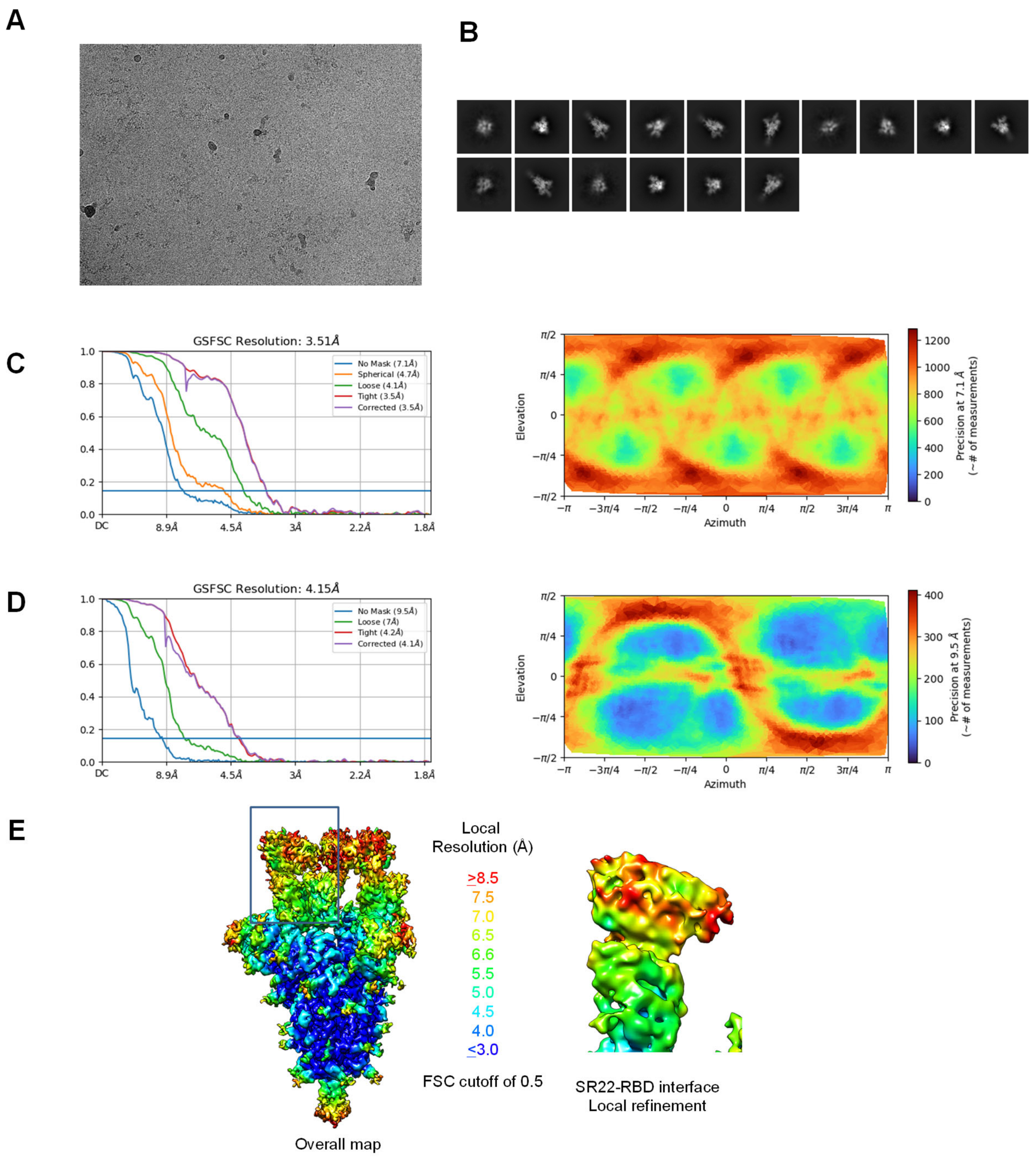


**Figure S7. Cryo-EM details of FSR22 in complex with SARS-CoV-2 S6P spike.** (**A**) Representative micrograph. (**B**) Representative cryo-EM 2D classes. (**C**) The Fourier shell correlation resulted in a resolution of 3.51 Å for the overall map using homogeneous refinement with C3 symmetry (left panel); the orientations of all particles used in the final refinement are shown as a heatmap (right panel). (**D**) The Fourier shell correlation resulted in a resolution of 4.15 Å for the masked local refinement of the RBD:SR22 interface (left panel) obtained using particle subtraction followed by local refinement; the orientations of all particles used in the local refinement are shown as a heatmap (right panel). **(E)** The local resolution of the final overall map and locally refined map is shown contoured at 4.5s and 8.1s, respectively. Resolution estimation was generated through cryoSPARC using an FSC threshold of 0.5.

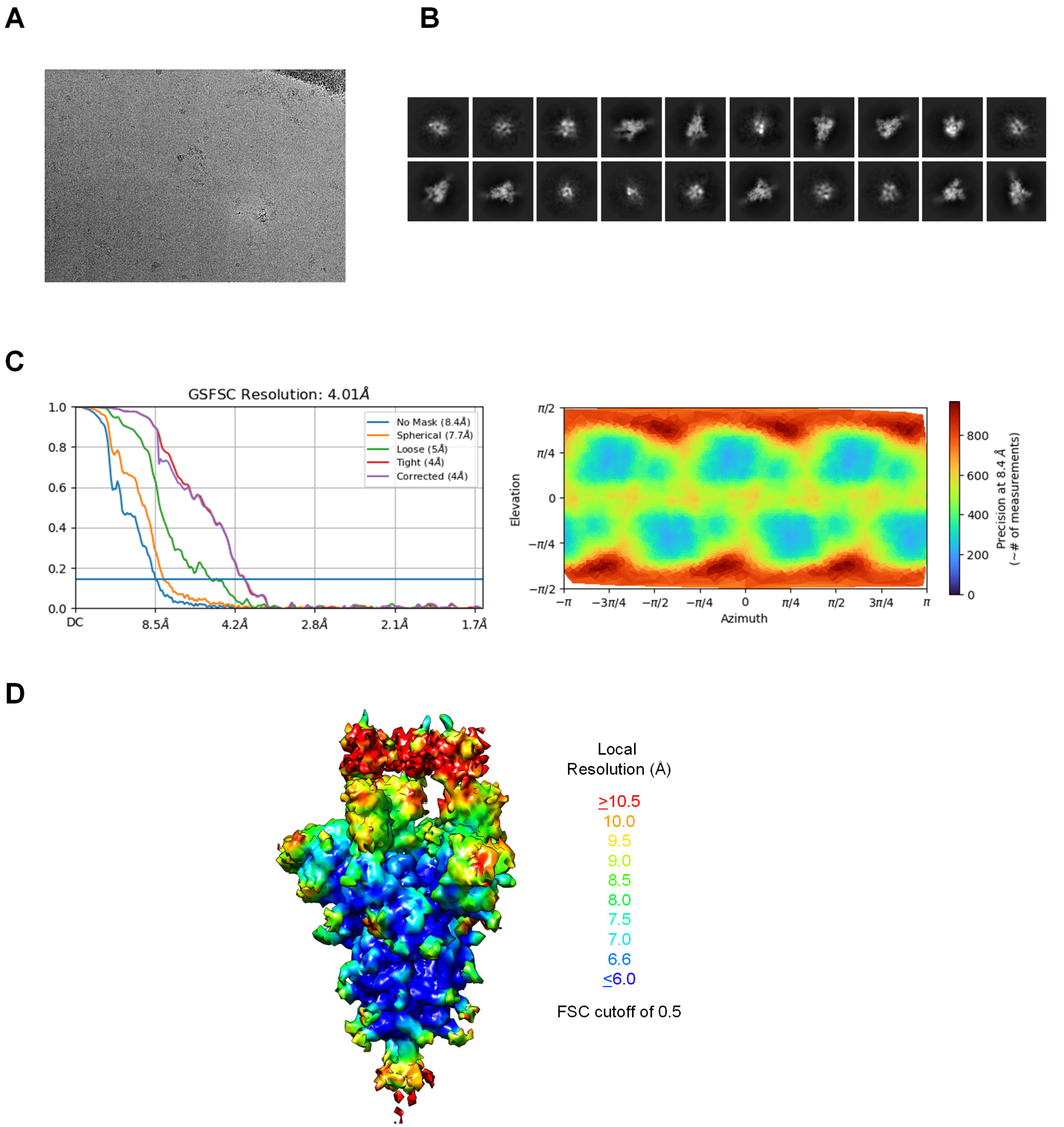


**Figure S8. Cryo-EM details of FSR16m in complex with SARS-CoV-2 S6P spike.** (**A**) Representative micrograph. (**B**) Representative cryo-EM 2D classes. (**C**) The Fourier shell correlation resulted in a resolution of 4.01 Å for the overall map using homogeneous refinement with C3 symmetry (left panel); the orientations of all particles used in the final refinement are shown as a heatmap (right panel). (**D**) The local resolution of the final overall map and locally refined map is shown contoured at 3.2σ. Resolution estimation was generated through cryoSPARC using an FSC threshold of 0.5.

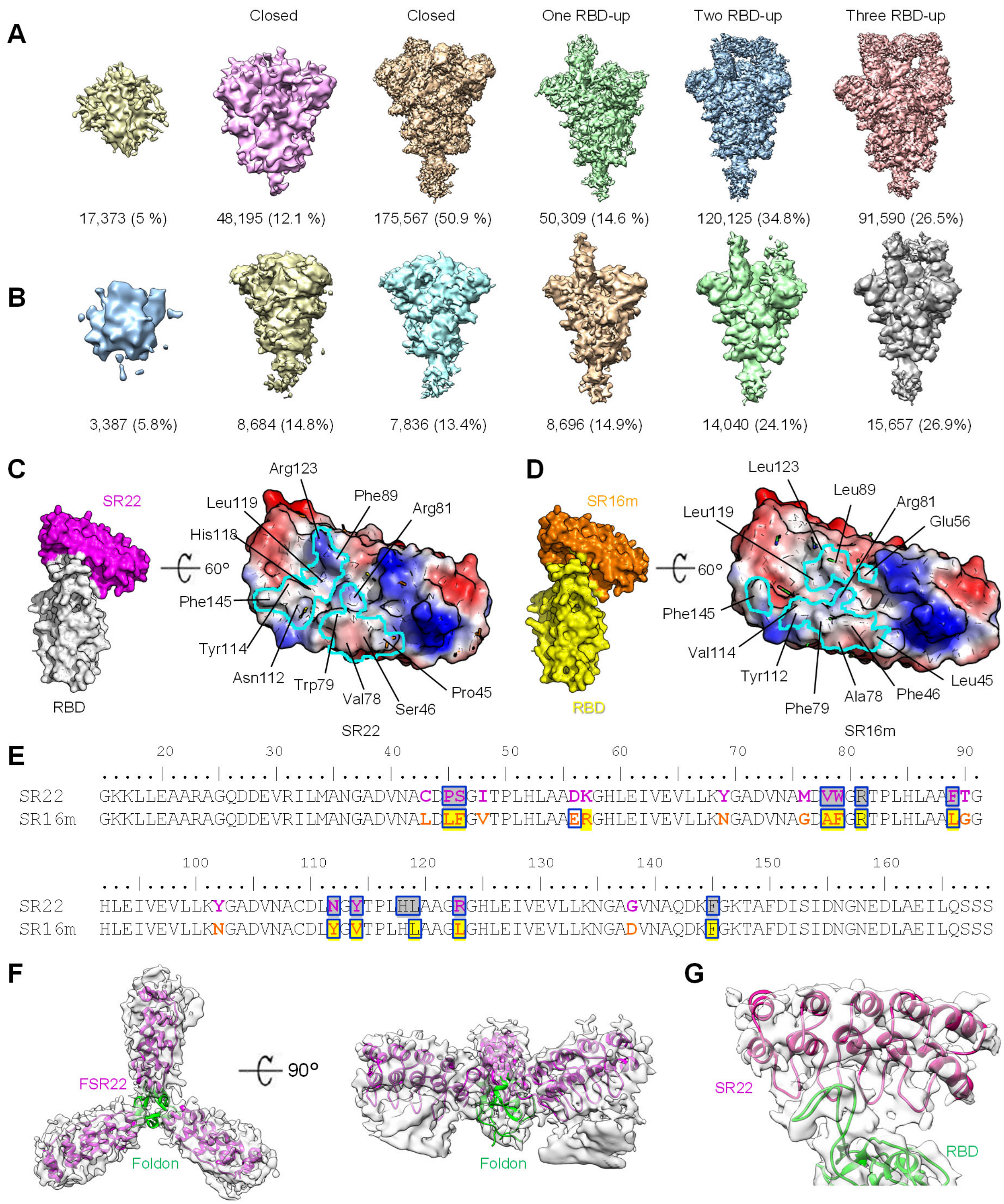


**Figure S9. FSR22 and FSR16m Structures.** (**A**) 3D-classes obtained from FSR22:SARS-CoV-2 spike complex. (**B**) 3D classes obtained from FSR16m:SARS-CoV-2 spike complex. (**C**) SR22:RBD complex (left). Electrostatic potential surfaces of SR22 (right). The footprint of RBD on SR22 is drawn in cyan. SR22 residues contacting RBD are shown (right). **(D)** SR16m:RBD complex (left). Electrostatic potential surfaces of SR16m (right). The footprint of RBD on SR16m is drawn in cyan. SR16m resides contacting RBD are shown (right). (**E**) Sequence alignment of SR22 and SR16m. Residues contacting RBD were shown in a rectangular box. (**F**) Three SR22s connected by foldon (green), was shown in electron density. (**G**) Electron density of SR22 and SR22-RBD interface.

A

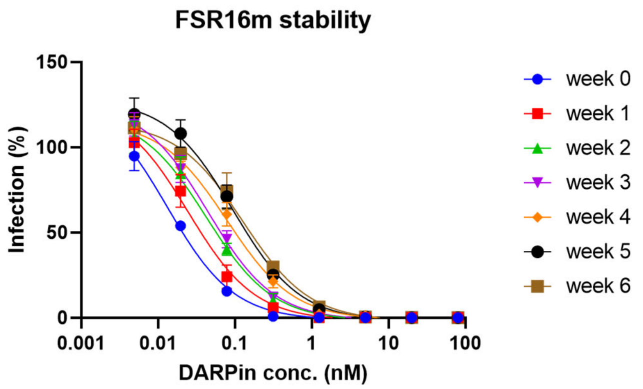


| | IC50 (nM) B.1.351 |
|---|---|
| Week 0 | 0.013 |
| Week 1 | 0.025 |
| Week 2 | 0.042 |
| Week 3 | 0.044 |
| Week 4 | 0.079 |
| Week 5 | 0.094 |
| Week 6 | 0.127 |

B

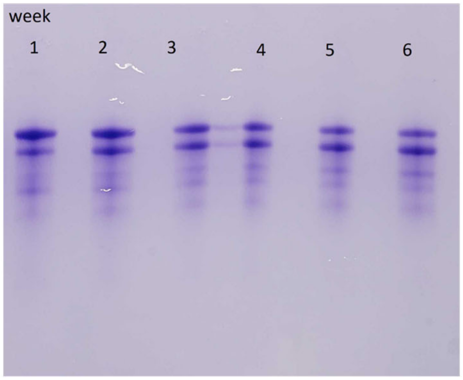


**Figure S10. Stability study.** FSR16m (0.85mg/ml, 11.87uM) was stored in PBS at room temperature. An aliquot was withdrawn each week and stored at -80˚C until the end of the study during which time the neutralization activity of all the aliquots were assayed using against lentivirus pseudotyped with spike protein from B.1.351 **(A)**. The same samples were also analyzed on a 12% Native PAGE gel. Due to the presence of a surface exposed cysteine residue, a small percentage of the protein formed disulfide bond during the storage in PBS.

### Table S1. Cryo-EM Data Collection and Refinement Statistics

| | SARS-CoV-2_ 6P in complex with FSR22 | SR22-RBD complex |
|---|---|---|
| **EMDB ID** | EMD-26200 | EMD-26201 |
| **PDB ID** | 7TYZ | 7TZ0 |
| Data Collection | | |
| Microscope | FEI Titan Krios | |
| Voltage (kV) | 300 | |
| Electron dose ($e^-/Å^2$) | 40 | |
| Detector | Gatan K3 BioQuantum | |
| Pixel Size (Å) | 0.873 | |
| Defocus Range (μm) | -1.0/-2.5 | |
| Magnification | 105,000 | |
| Reconstruction | | |
| Software | cryoSPARC v3.3 | |
| Number of particles extracted | 417,462 | |
| Number of particles used | 18,892 | |
| Symmetry | C3 | C1 |
| Box size (pix) | 500 | 500 |
| Resolution (Å) ($FSC_{0.143}$) | 3.5 | 4.2 |
| Refinement | | |
| Software | Phenix 1.20 | |
| Protein residues | 3,711 | 347 |
| CC (mask) | 0.79 | 0.62 |
| R.m.s. deviations | | |
| Bond lengths (Å) | 0.003 | 0.003 |
| Bond angles (°) | 0.561 | 0.599 |
| Validation | | |
| Molprobity score | 1.94 | 2.02 |
| Clash score | 7.89 | 10.4 |
| Favored rotamers (%) | 99.7 | 100 |
| Ramachandran | | |
| Favored regions (%) | 91.3 | 92.4 |
| Allowed regions (%) | 8.68 | 7.58 |
| Disallowed regions (%) | 0.03 | 0 |

14. Case JB, Mackin S, Errico J, Chong Z, Madden EA, Guarino B, Schmid MA, Rosenthal K, Ren K, Jung A, Droit L, Handley SA, Halfmann PJ, Kawaoka Y, Crowe JE, Fremont DH, Virgin HW, Loo Y-M, Esser MT, Purcell LA, Corti D, Diamond MS. Resilience of S309 and AZD7442 monoclonal antibody treatments against infection by SARS-CoV-2 Omicron lineage strains. bioRxiv. 2022:2022.03.17.484787. doi: 10.1101/2022.03.17.484787.
15. Case JB, Bailey AL, Kim AS, Chen RE, Diamond MS. Growth, detection, quantification, and inactivation of SARS-CoV-2. Virology. 2020;548:39-48. doi: 10.1016/j.virol.2020.05.015.
16. Hassan AO, Case JB, Winkler ES, Thackray LB, Kafai NM, Bailey AL, McCune BT, Fox JM, Chen RE, Alsoussi WB, Turner JS, Schmitz AJ, Lei T, Shrihari S, Keeler SP, Fremont DH, Greco S, McCray PB, Jr., Perlman S, Holtzman MJ, Ellebedy AH, Diamond MS. A SARS-CoV-2 Infection Model in Mice Demonstrates Protection by Neutralizing Antibodies. Cell. 2020;182(3):744-53 e4. Epub 2020/06/20. doi: 10.1016/j.cell.2020.06.011. PubMed PMID: 32553273; PMCID: PMC7284254.
17. Vanblargan L, Adams L, Liu Z, Chen RE, Gilchuk P, Raju S, Smith B, Zhao H, Case JB, Winkler ES, Whitener B, Droit L, Aziati I, Shi P-Y, Creanga A, Pegu A, Handley S, Wang D, Boon A, Crowe JE, Whelan SPJ, Fremont D, Diamond M. A potently neutralizing anti-SARS-CoV-2 antibody inhibits variants of concern by binding a highly conserved epitope. 2021.
18. Olia AS, Tsybovsky Y, Chen SJ, Liu C, Nazzari AF, Ou L, Wang L, Kong WP, Leung K, Liu T, Stephens T, Teng IT, Wang S, Yang ES, Zhang B, Zhang Y, Zhou T, Mascola JR, Kwong PD. SARS-CoV-2 S2P spike ages through distinct states with altered immunogenicity. J Biol Chem. 2021;297(4):101127. doi: 10.1016/j.jbc.2021.101127. PubMed PMID: 34461095; PMCID: PMC8393506.
19. Punjani A, Rubinstein JL, Fleet DJ, Brubaker MA. cryoSPARC: algorithms for rapid unsupervised cryo-EM structure determination. Nature Methods. 2017;14(3):290-6. doi: 10.1038/nmeth.4169.
20. Benton DJ, Wrobel AG, Roustan C, Borg A, Xu P, Martin SR, Rosenthal PB, Skehel JJ, Gamblin SJ. The effect of the D614G substitution on the structure of the spike glycoprotein of SARS-CoV-2. Proceedings of the National Academy of Sciences. 2021;118(9):e2022586118. doi: 10.1073/pnas.2022586118.
21. Jumper J, Evans R, Pritzel A, Green T, Figurnov M, Ronneberger O, Tunyasuvunakool K, Bates R, Žídek A, Potapenko A, Bridgland A, Meyer C, Kohl SAA, Ballard AJ, Cowie A, Romera-Paredes B, Nikolov S, Jain R, Adler J, Back T, Petersen S, Reiman D, Clancy E, Zielinski M, Steinegger M, Pacholska M, Berghammer T, Bodenstein S, Silver D, Vinyals O, Senior AW, Kavukcuoglu K, Kohli P, Hassabis D. Highly accurate protein structure prediction with AlphaFold. Nature. 2021;596(7873):583-9. doi: 10.1038/s41586-021-03819-2.
22. Emsley P, Cowtan K. Coot: model-building tools for molecular graphics. Acta crystallographica Section D, Biological crystallography. 2004;60(Pt 12 Pt 1):2126-32. Epub 2004/12/02. doi: 10.1107/s0907444904019158. PubMed PMID: 15572765.
23. Adams PD, Afonine PV, Bunkóczi G, Chen VB, Davis IW, Echols N, Headd JJ, Hung L-W, Kapral GJ, Grosse-Kunstleve RW, McCoy AJ, Moriarty NW, Oeffner R, Read RJ, Richardson DC, Richardson JS, Terwilliger TC, Zwart PH. PHENIX: a comprehensive Python-based system for macromolecular structure solution. Acta crystallographica Section D, Biological crystallography. 2010;66(Pt 2):213-21. Epub 2010/01/22. doi: 10.1107/S0907444909052925. PubMed PMID: 20124702.
24. Williams CJ, Headd JJ, Moriarty NW, Prisant MG, Videau LL, Deis LN, Verma V, Keedy DA, Hintze BJ, Chen VB, Jain S, Lewis SM, Arendall WB, 3rd, Snoeyink J, Adams PD, Lovell SC, Richardson JS, Richardson DC. MolProbity: More and better reference data for improved all-atom structure validation. Protein science : a publication of the Protein Society. 2018;27(1):293-315. Epub 2017/11/27. doi: 10.1002/pro.3330. PubMed PMID: 29067766.
25. Barad BA, Echols N, Wang RY, Cheng Y, DiMaio F, Adams PD, Fraser JS. EMRinger: side chain-directed model and map validation for 3D cryo-electron microscopy. Nat Methods. 2015;12(10):943-6. Epub 2015/08/19. doi: 10.1038/nmeth.3541. PubMed PMID: 26280328; PMCID: PMC4589481.
26. Pettersen EF, Goddard TD, Huang CC, Meng EC, Couch GS, Croll TI, Morris JH, Ferrin TE. UCSF ChimeraX: Structure visualization for researchers, educators, and developers. Protein Sci. 2021;30(1):70-82. Epub 2020/09/04. doi: 10.1002/pro.3943. PubMed PMID: 32881101; PMCID: PMC7737788.
